# Skin photoreceptors in the leopard frog

**DOI:** 10.1101/161364

**Authors:** George Wald, Stephen Rayport

## INTRODUCTION

It is an engaging thought that the great sense organs concentrated in the head region of vertebrates were all derived from primitive systems of widely diffused skin receptors. In this sense, one can think of the receptors of hearing as derived from cutaneous touch spots, with which they share many physiological properties; the organ of equilibrium the cristae of the semicircular canals from deep pressure receptors; smell from chemoreceptors that remain widely distributed over the bodies of the mucous - skinned fishes and amphibia and have withdrawn into the mucous area of the mouth in land vertebrates. In all these cases, the transition from skin receptors to sense organ has involved four parallel developments: 1) The concentration of diffuse systems of individual end organs into a sensory tissue — the basilar membrane of the ear, retina of the eye, the cristae of the semicircular canals, the olfactory patch. 2) The elaboration of accessory structures that concentrate the appropriate stimulus upon the sensory surface and exclude other stimuli: the eye, ear and nose. 3) The afferent nerve nuclei, which in the case of the skin receptors lie close to the spinal cord in the dorsal root ganglia have migrated out into the sense organ itself, so that the sensory tissue is virtually an outpocket of the brain, able to process as well as respond to sensory stimuli. 4) The perceptions associated with the great sense organs display the extraordinary feature of projection to a distance. Whereas sensations from skin receptors are perceived locally, more or less where the stimulus acts (hence proprioceptors), the sensations from the great sense organs are perceived as though projected to where the stimulus originates (hence distance receptors, exteroreceptors). No amount of introspection lets us smell odors at the olfactory patch, hear tones in the cochlea, or see images on our retinas. Yet of course these are the places where the stimuli act, much more deeply buried than the skin receptors, yet always perceived as at a distance.

Thought of in this way, what should be the skin analogs of the visual organs? Hot spots, since they can respond to infrared radiation, indeed overlapping in this regard with the human retina (Griffin et al., 1947)? The infrared organ of pit vipers would represent a more acceptable analog. Time and again skin photoreceptors have been reported in man, yet always under highly questionable circumstances and never surviving careful examination.

For these reasons the allegations that blinded frogs orient to light (Parker, 1903) and that blinded tadpoles are stimulated by light to swim (Obreshkove, 1921) are of great interest. Moreover, Pearse (1910) and Laurens (1911) report a definite preference of various intact and eyeless amphibians for blue lights in a series of behavioral studies. More recently, Steven studied the dermal light sense of two other vertebrates, the brook lamprey (Steven, 1950) and the hag (Steven, 1955). In both cases, a spectral sensitivity was found maximal at about 520 nanometers (nm) in the brook lamprey and 510 nm in the hag. These agree well with Wald’s finding of porphyropsin (absorption maximal at about 522 nm) in fresh water animals, and rhodopsin (maximum at about 500 nm) in ocean and land animals (Wald, 1945, 1960, 1963).

Electrophysiological evidence pointing to the existence of skin photoreceptors in amphibians comes from studies by Becker with Cone in 1966 (1966) and with Goldsmith in 1968 (Becker and Goldsmith, 1968) and 1970 (Becker, 1970). In frog skin, Becker and Cone (1966) found a fast potential, which we term the fast skin photo - voltage (FSP), that is remarkably similar to the pigment epithelium component of the electroretinogram, and a late skin response, which we call the electrodermogram (EDG), which followed the early response with a latency of about three quarters of a second at room temperature. This monophasic response lasted as much as five to ten seconds, taking one to two seconds to rise to a maximum and from four to eight seconds to return to the base line (see Fig. 3). Over the entire intensity range available, Becker and Cone found that both components of the light - evoked skin response were linear with intensity. Neither showed any adaptation to repeated stimuli. The FSP was indeed mediated by a black pigment, presumably melanin, as all three components of the FSP (the alpha, beta and gamma waves) showed a uniform, flat spectral sensitivity. On the contrary, the late response had a spectral sensitivity rising to a maximum in the violet or near - ultraviolet, and was relatively insensitive to light of wavelength longer than about 600 nm. Becker (1970) suggests that the late response is evidence of the discharge of skin glands in response to light stimulation. He supports this idea with two points: 1) The effect of adrenalin, which significantly enhances the late response, is blocked by theophylline, a caffeine analog; presumably, adrenalin sensitizes the glands to light while theophylline relaxes the glandular smooth muscle. 2) The late response is larger when light stimulation is to the inside rather than the outside of the skin, the glands being beneath the pigmented layers of the skin.

**FIGURE 3.**
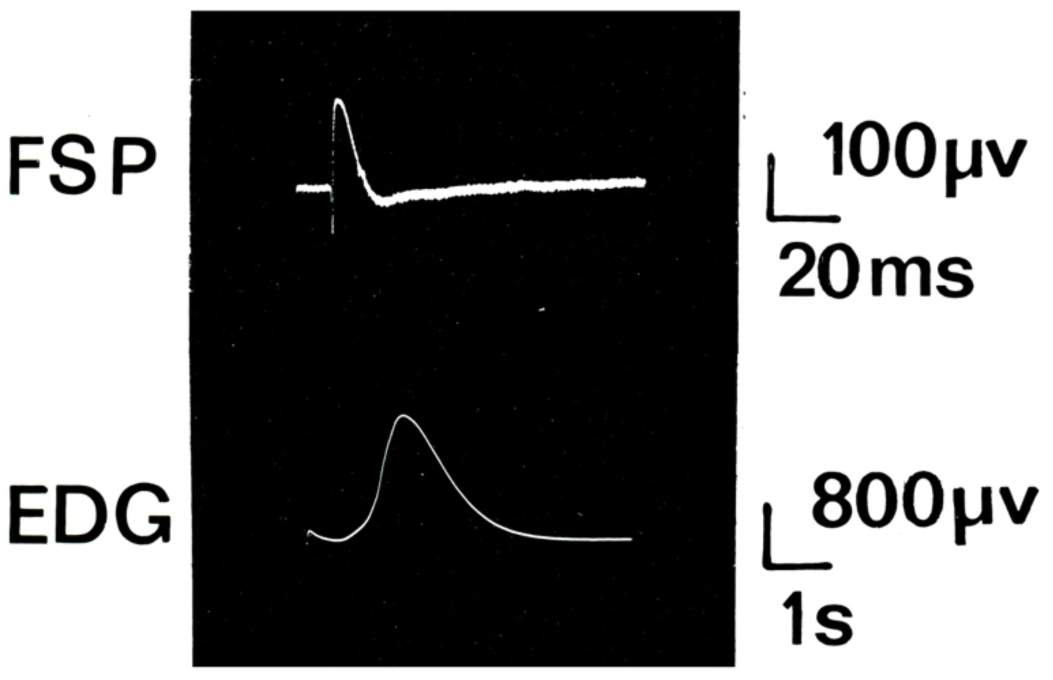
Light-evoked electrical responses in frog skin have two components. The fast component, the fast skin photo-voltage (FSP), is triphasic. The alpha wave which has no measureable latency is followed by the beta and gamma waves. The gamma wave lasts until or past the start of the later slow response (with capacity coupled amplification, the gamma wave appears to be over after 50 to 100 msec), the electrodermogram (EDG). The bottom trace, an expansion of the second response, shows that the EDG has a latency of about 0.75 sec at 23 °C (the end of the gamma wave distorts the beginning of the EDG). Upward deflections correspond to hyperpolarization of the skin (outside becomes more negative). The baseline voltage on the latter two traces is the same as the minimum voltage immediately before the EDG. The resting skin potential is at about 15 millivolts.

It is important to know that frog skin maintains a potential difference of as much as 100 mv, outside negative. The potential is created by the active transport of sodium from the outside to the inside medium (Ussing and Zerahn, 1951). Coupled to the inward sodium transport is a potassium transport from the inside medium into the epithelial cells from which the potassium leaks back into the inside medium (Ussing, 1960; Macrobbie and Ussing, 1961). Chloride is carried from the outside to the inside medium at a rate about 2 to 5% that of sodium transport (Kristensen, 1972), so that to a good approximation, the skin potential is the result exclusively of an inward sodium current.

Variations in the skin potential could be mediated by modulation of any of the above permeability and transport phenomena. In addition, the skin glands cause a variation of the skin potential when they discharge (Schoffeniels and Salee, 1965; Lindley, 1969). Apparently, the contents of the glands are chloride - rich, thus creating a hyperpolarization when they pulse their contents to the outside. When the skin glands open, a reverse (depolarizing) current develops due to the large surface area exposed (about a third that of the skin (Lindley, 1969). Also the cells lining the glands and especially the ducts are known to take up chloride, further adding to the current (Lindley, 1969).

Frog skin consists of epidermal and dermal layers. The epidermis is the active physiological portion of the skin, while the dermis serves structural and supportive functions. All the electrical activity of interest here takes place in the epidermis and in the outermost layer of the dermis where the skin glands are found. The skin potential is entirely across the epidermis. *Zonulae occludentes* in the outer layers of the epidermis, as well as other tight junctions, subserve crucial insulating and structural functions (Farquhar and Palade, 1965).

Ultrastructural investigations of frog skin in search of skin photoreceptors have been inconclusive. We found stacked membranes (Fig. 1) which however seem to correlate with pterinosomes (non - melanophore pigment granules) which Matsumoto found in the skin of the red - scaled swordtail, *Xiphophorus* (see Bagnara, 1966). Nakao (1974) has reported the discovery of lamellated bodies in the space between epidermis and dermis of the frog tadpole tail in *Rana rugosa*. These bodies are on the order of a twentieth of a micron in diameter, with lamina of 20 Å thickness separated by 20 to 30 Å gaps. The structures thus have about ten layers and are continuously distributed throughout the area studied. Crucial to this study was the use of phosphotungstic acid, as these lamellated bodies were never found when it was omitted in the preparation of the tissue. To our knowledge, there are no other ultrastructural reports suggestive of skin photoreceptors.

**FIGURE 1.**
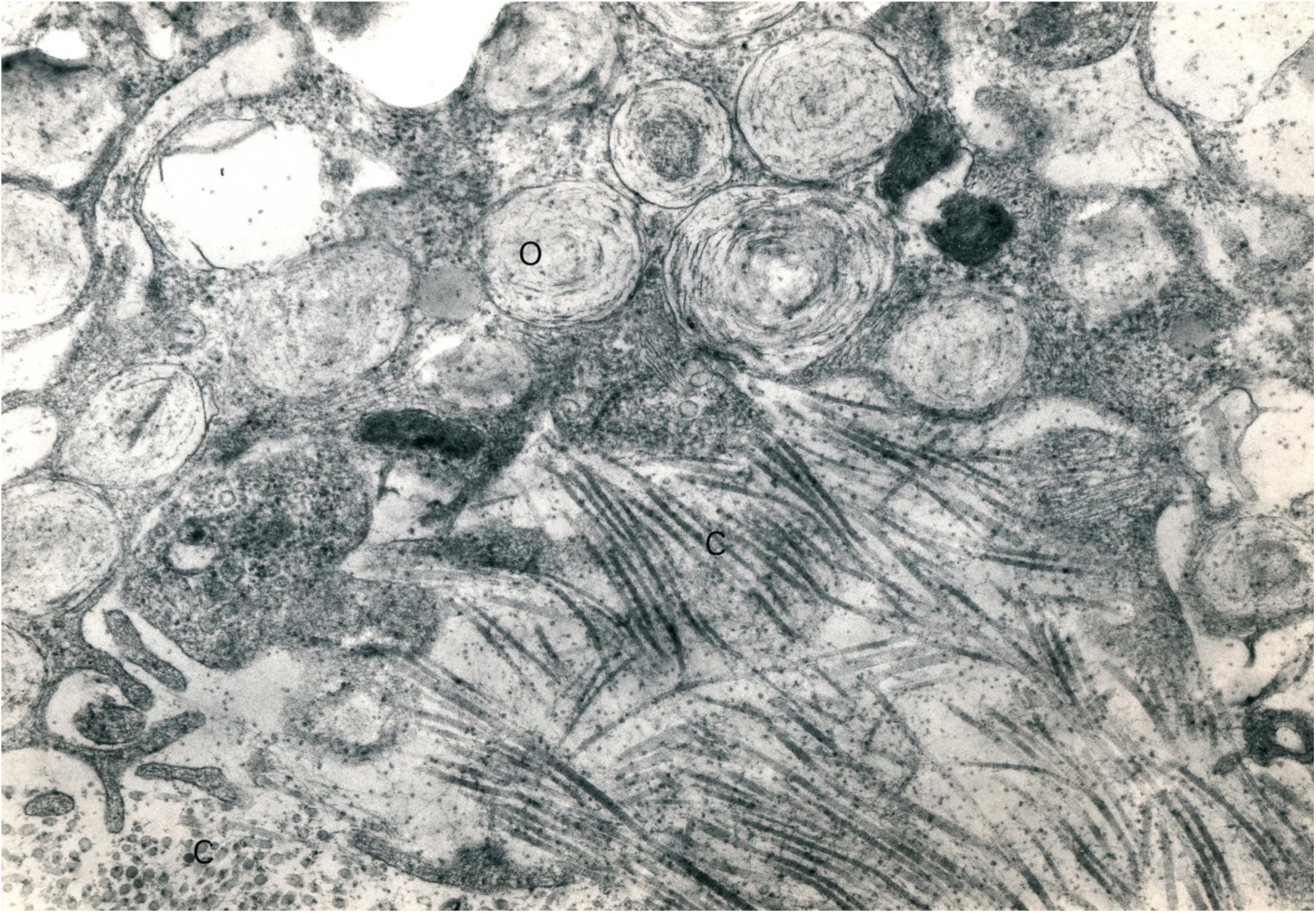
Section through the dermis at the junction between the stratum spongiosum and the compactum. Specimen from the dorsal skin of *Rana pipiens* showing the structures we have called ‘onions’ (0) which are identified as non - melanin containing pigment granules. They are very plentiful in this layer. Collagen fibers are seen in two orientations at right angles to each other, longitudinal (C) and transverse (C). Tissues fixed in glutaraldehyde and post-fixed in osmic acid. X32,000

The electrophysiological study of receptors in frog skin dates from the work of E.D. Adrian (1928). The skin nerves are admirably suited for study because they are both easily accessible and have few enough fibers that one can distinguish the activity of single fibers (Spray, 1974a; Sutherland and Nunnemacher, 1974). In short, tactile receptors are free nerve endings in the epidermis. Deep pressure and general chemoreceptors are bulbous nerve endings in the outer layer of the dermis, the stratum spongiosum (Rubin and Syrocki, 1936). These deeper receptors also may include cold receptors (Spray, 1974b).

We have found evidence strongly suggesting that there are indeed frog skin photoreceptors. The receptors have a rhodopsin - like spectral sensitivity curve, they undergo adaptation and photorecovery indicating that there is an interconvertible pigment with at least two states. We suggest a mechanism for the EDG. But both behavioral significance and thus function remain elusive. This work was published initially in abstract form (Wald and Rayport, 1974).

## EXPERIMENTAL METHODS

The electrical characteristics of frog skin were measured utilizing a modified Ussing cell (Ussing and Zerahn, 1951) **(Fig. 2)**. A circle of skin of 0.13 cm^2^ was exposed. It separated two 100 ml baths of Ringer’s Solution (NaCl 110 *mM*, KCl 2.5 *mM, C*aCl_2_ 2.5 mM buffered to pH 7.4) into which dipped light - shielded, non - polarizable, silver - silver chloride electrodes. The Ringer’s solution was continuously oxygenated by a bubble lift in both chambers.

**FIGURE 2.**
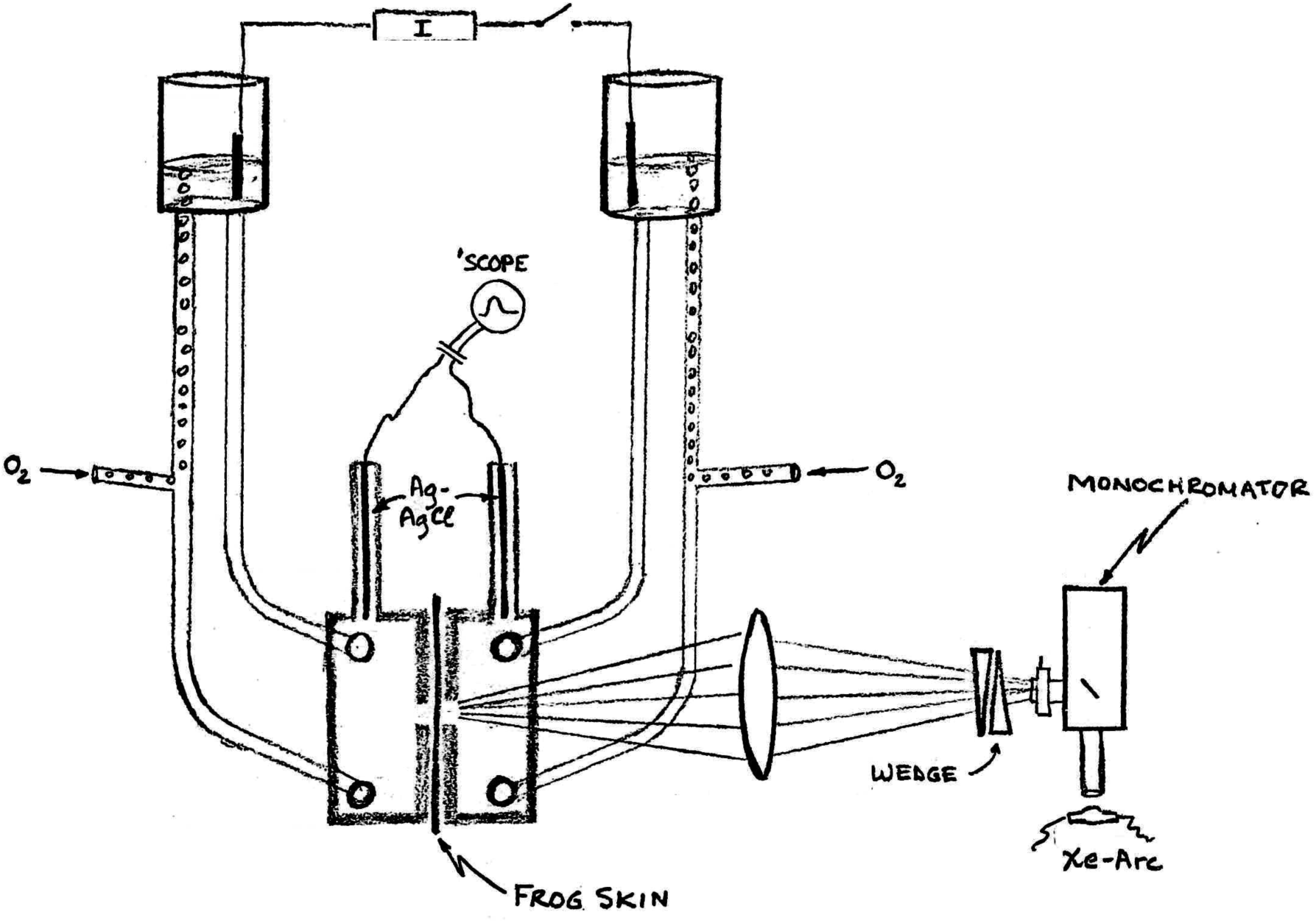
Ussing cell apparatus. Frog skin is placed in Ussing cell between two chambers, bubbled with oxygen (O_2_), circulated with the bubble lift. Continuous current (I) is applied via silver - silver chloride (Ag - AgCl) electrodes in the fluid reservoirs; the skin potential is measured with a second set of electrodes in the Ussing cell. Voltages are displayed and photographed from an oscilloscope (‘SCOPE). Light stimulation is delivered via a monochromator, with a Xenon - arc source (Xe - Arc); the intensity is regulated with neutral density filters (WEDGE) and focused on the skin with a lens (oval).

Areas of skin were taken routinely from the backs of the animals. In the case of the leopard frog, the skin exposed was a non – black spot. Light stimulation impinged on the outer surface of the skin. In view of the intensities of light involved, it was considered advisable to minimize possible heating effects. The maximum intensity used was 10^18^ quanta *I* cm^2^ sec.

Light from a 150 watt Osram Xenon arc lamp was projected through a 250 mm grating monochromator (Bausch & Lomb, grating 1200 lines per mm), with its slits set at 3 mm giving an effective bandwidth of about 10 nm. A photographic shutter at the exit slit determined the stimulus duration, and a lens projected the image of the exit slit onto a pair of circular neutral density wedges, rotating in opposite directions so as to compensate each other. The wedge was calibrated using a Welch Densichron (Welch Scientific Co.). A range of densities of about 3.5 orders of magnitude was available. Finally, the light beam exiting from the wedge was focused down to a 4 mm square where the skin was placed. The optical system was all quartz including the monochromator and the windows of the Ussing cell (Fig. 2).

The system was calibrated by placing a Reeder thermocouple at the location of the skin. The thermocouple voltage was amplified by a Brower laboratories Model 261 preamplifier connected to their Model 131 Lock - in - Voltmeter. The light signal was chopped at 13 Hz and the rectified voltage read off the Model 131. After correction for heat filters (used to limit the calibration to the visible spectrum) and energy per photon, the relative energy distribution was obtained.

The Ussing cell and the flow system were shielded in a grounded copper cage. The skin potential recorded from the silver - silver chloride electrodes was amplified with a Grass P - 18 DC - preamplifier (Bandwidth 0 - 100 Hz). The amplified signal was displayed on Tektronics 502A and 565 oscilloscopes from which the records were photographed with a Polaroid camera.

Current passed through the skin to adjust the skin potential was applied through a second set of silver - silver chloride electrodes in the fluid reservoirs. The output of a Grass S - 4 Stimulator, on the constant voltage setting, was passed through a Stimulus Isolation Unit (Grass) and two 100K ohm resistors (one at each terminal of the output) in series, with the preparation. Since the skin resistance in the chamber was about 2,000 ohms, this made an excellent current source. Current was monitored with a series microamplifier; 100 microamps was the maximum current passed.

*Rana pipiens* were obtained from various suppliers. Sigma chemicals were used.

## OBSERVATIONS

In the frog, light stimuli to the skin evoke a modulation of the trans - skin potential. These variations of the skin potential have differing time courses and spectral sensitivities. The early response or fast skin potential (FSP) consists of alpha, beta and gamma waves which appear in characteristic order (Fig. 3). We, as Becker et al. (Becker and Cone, 1966; Becker and Goldsmith, 1968; Becker, 1970) have found that all the components of the FSP are equally sensitive to all wavelengths of the visible spectrum. On the contrary, the monophasic late response, which we term the electrodermogram (EDG), is maximally excited by near - ultraviolet light. Repeated stimulation in the near - ultraviolet gives a diminishing response which reaches a steady state at an amplitude many times lower than the original maximum. Yellow light reverses the adaptation and facilitates the EDG.

The latency of the EDG is about 0.75 sec at 23° C (Fig. 3). It takes about one sec to peak, and about another two to three secs to return to the baseline. At skin potentials above 50 mv outside negative, light stimuli generate a depolarizing EDG. Below 50 mv trans skin potential, the response is hyperpolarizing (Fig. 4A). EDG responses of comparable amplitudes at skin potentials on different sides of the 50 mv reversal potential, when superimposed with one inverted (Fig. 4B), show slightly different time courses and wave shapes.

**FIGURE 4.**
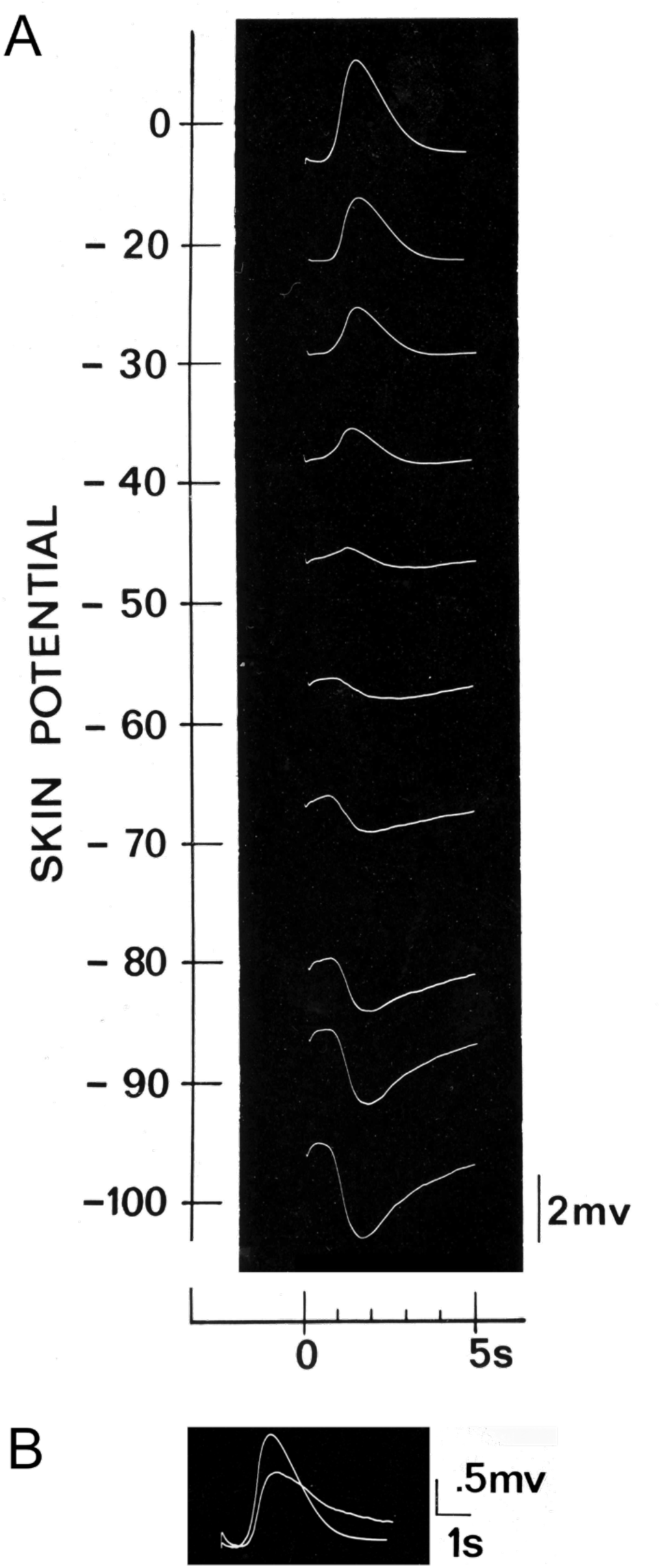
EDG depends trans-skin potential. A Setting the skin potential by passing current externally allows evaluation of the dependence of the EDG on skin potential. Below the critical 50 mv potential (outside negative), the EDG response is hyperpolarizing; above 50 mv it is depolarizing. Reversing the skin potential to outside positive does not increase the EDG beyond its amplitude at zero potential. On the contrary, with increasingly greater negative skin potentials, the EDG amplitude increases proportionally. B) The EDG has opposite polarities at 0 and 100 mv skin potential, as well as a slightly, different latency, amplitude, and delay in returning to the base - line voltage. The larger amplitude response is at 0 mv skin potential and the smaller at 100 mv outside negative.

Up to the maximum intensity used 10^18^ quanta / cm sec^2^ the EDG amplitude is linear with intensity (Fig. 5). However, this is not the case for the variation of the EDG amplitude with stimulus duration. A plot of EDG amplitude with respect to duration is sigmoidal (Fig. 5). The linearity of EDG amplitude with stimulus intensity simplified the determination of the spectral sensitivity.

**FIGURE 5.**
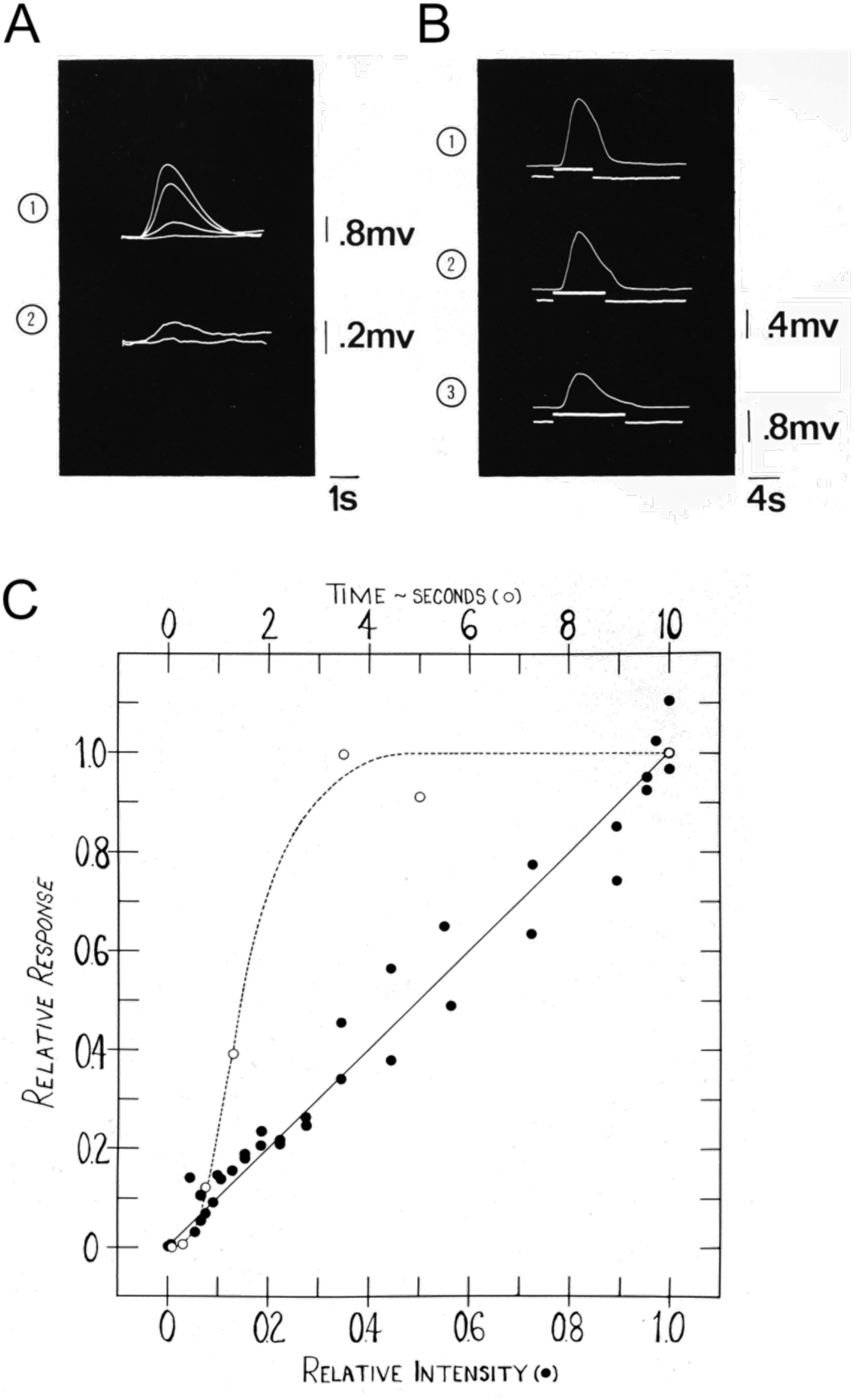
EDG as a function of stimulus intensity and duration. Stimulus intensity and duration effects. A) EDG in response to light stimuli of different intensities. Responses to lights of relative intensities of 0, 0.1, 0.5, 0.6 log units. Responses to lights of relative intensities of 0.6, 1.0 log units. The latency of the EDG is the same at all of these intensities, but the latency to peak diminishes with intensity. B) EDG in response to long duration stimuli: 5 sec, 7 sec, 10 sec. The skin potential here is about 15 mv. At higher potentials, the response increases in amplitude after the initial peak, rather than decreasing as is seen here. C) The EDG amplitude is linear with stimulus intensity to the limits of the stimuli available (about 5^12^ quanta / cm^2^ sec). The EDG amplitude is sigmoidal with respect to duration of stimulus. Effective stimulation at shorter times than 0.1 sec has been the exception, and after two to four seconds, the EDG peaks before the stimulus is ended so that continuous stimuli cannot produce a larger EDG response.

To determine the spectral sensitivity, a saturating yellow recovery light was presented followed by a test stimulus. A control test stimulus was presented alternately to assure standardization of the measurements. Relative sensitivity was determined by scaling by the quanta / sec at each wavelength. Fig. 7 shows the spectral sensitivity of the EDG on an equal energy basis. The spectral sensitivity curve has a maximum at about 385 nm, and a half - amplitude bandwidth of about 83 nm. Effectively. no stimulation occurs below 310 nm, nor above 500 nm.

**FIGURE 7.**
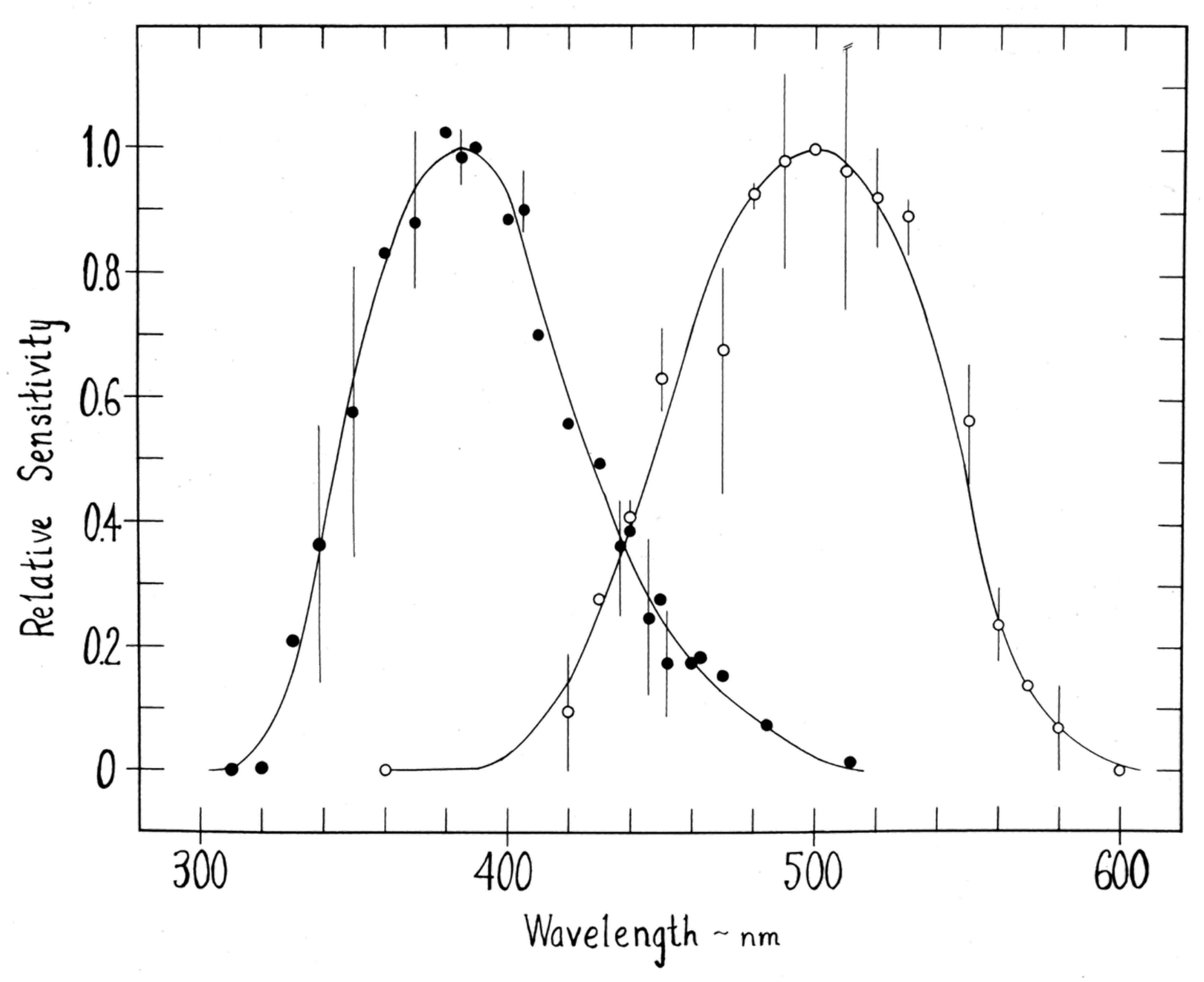
Spectral sensitivities of frog skin photoreceptors. Excitation is maximally sensitive at about 385 nanometers (filled circles). Recovery is maximal at about 500 nm (open circles). The excitation curve has a half - amplitude bandwidth of about 83 nm and the recovery curve a bandwidth of 101 nm. Points with error bars represent the mean of two to four measurements in different frogs. The error bars extend from the maximum to the minimum response observed at the particular wavelength. The standard wavelength used to correct the raw data for each curve was the maximum wavelength (385 and 500 nm).

The EDG adapts to repeated stimulation. Yellow light although itself causing no change in the skin potential, restored the EDG amplitude to its original or greater amplitude (if it had been partially adapted to start with). In Fig. 6 the phenomenon is illustrated using broad - bandpass filters. First, successive stimuli through a Corning 9863 filter (transmission maximum at 330 nm, range from 235 nm to 400 nm) are superimposed, showing the diminishing amplitude of the EDG. Then, a single flash through a Corning 3385 filter (transmission greater than 470 nm) is presented, giving no measurable response. But when the C9863 filter is subsequently used, the EDG has recovered and can be adapted once again with repeated flashes.

**FIGURE 6.**
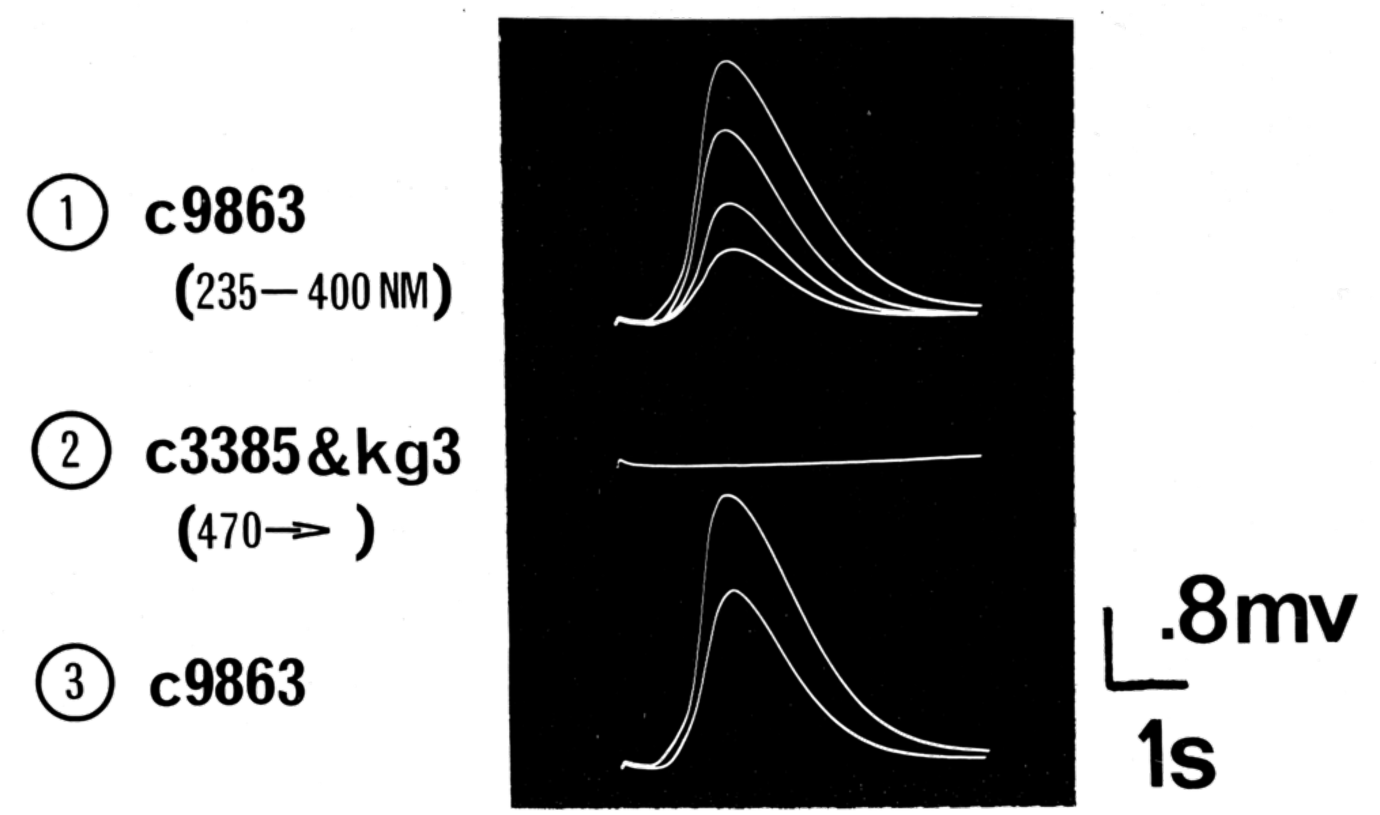
Adaptation - recovery phenomenon in skin photoreceptors. Successive excitatory stimulation produces adaptation, while a recovery stimulus which produces no excitation regenerates the EDG amplitude. Here, ① excitation is through a near - ultraviolet transmitting filter (C9863; 235 – 400 nm) which when presented at regular intervals of 30 sec produces significant adaptation. ② A single flash of a yellow transmitting filter (C3385; greater than 470nm, with KG3, a heat filter) photoregenerates the EDG response without itself producing any excitation. ③ The photo-regenerated EDG can again be adapted. This process has been repeated in some experiments for upwards of ten hours, several hundred times.

When the frog skin is left in an adapted state for as long as 70 min in the dark at 23 °C, no recovery is found, whereas as little as a fraction of a sec of yellow light following the hour’s wait can restore the response. The absence of dark recovery indicates, that at least on the scale of an hour, no dark reactions take place.

The relative effectiveness of different recovery stimuli was evaluated by calculating the net increase in the amplitude of the EDG in response to a near - ultraviolet stimulus. First this standard near ultra - violet light was presented for several seconds, adapting the EDG considerably. The same standard stimulus was then presented for a sec and the EDG amplitude recorded. A recovery test stimulus was presented and immediately after the response to the standard wavelength stimulus remeasured. The amount of recovery was taken to be the difference in EDG amplitude between post - and pre - recovery. To correct for variations in the skin sensitivity, a standard recovery stimulus was regularly presented and the test responses normalized. The spectral sensitivity of photo - recovery is maximal at about 500 nm, with a half - amplitude bandwidth of about 100 nm (Fig. 7). No recovery could be measured below about 400 nm, nor above 600 nm. The fact that the spectral sensitivity curves for both excitation and recovery overlap at least from 400 to 500 nm suggests that some correction is indicated (see Discussion).

Since we have measured the spectral sensitivities of excitation and recovery in different ways, comparing the sensitivities of the two systems is difficult. However, if alternating excitatory and recovery stimuli are properly balanced, then each excitatory stimulus should yield an invariant response. In such a steady state situation, the only unknown is the relative sensitivities of the two systems. Using the visual transmissions (T_v_, the ratio of the effect of the filtered stimulus to the unfiltered stimulus) of the excitatory and recovery stimuli, we find that the excitatory sensitivity is about twice that of the recovery sensitivity.

## DISCUSSION

The light - evoked electrical response from frog skin has two components, the fast skin photovoltage (FSP) and the slower electrodermogram (EDG). The EDG has been our prime focus. It is mediated by at least a two - state system; more than two states might be resolved at lower temperatures. Judging by the spectral sensitivities, stimulation and concurrent adaptation are mediated by a yellow pigment absorbing maximally at about 385 nm. The action of light causes a variation of the skin potential, associated with the conversion of 385 - pigment to 500 - pigment. The purple 500 - pigment is converted back to 385 - pigment with the recovery action of yellow light. The system sensitivity is then adapted when the level of 500 - pigment is high and restored when the level of 385 - pigment is high.

Our findings suggest a strong parallel between frog skin photoreceptors and invertebrate photoreceptor photochemistry (cf. Hamdorf et al., 1973). As in invertebrates, we have been studying a system with two states which can be described as follows:

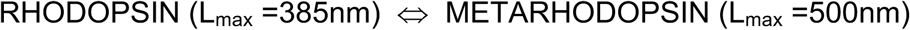

At room temperature, the system is always in equilibrium between the two states. As in invertebrates, dark interconversions must take place very slowly through renewal of the photoreceptor membranes and not by chemical regeneration; regeneration is exclusively mediated by light. In the overlap region of the frog skin receptor system, 400 to 500 nm, a monochromatic light simultaneously stimulates, adapts and recovers the system.

There are potential advantages of a system with large overlap over one with no overlap or one with complete overlap (as in the squid, *Loloigo pealii*; see Wald’s discussion of Hamdorf et al. (1973)). With no overlap, complete adaptation is probable and the system then becomes useless. In the case of total overlap, the photo - equilibrium between Rhodopsin and Metarhodoposin is independent of the wavelength of light stimulation and only dependent on the quantal efficiencies of the rates of interconversion between the two states, so that the system has only monochromatic capabilities. In the intermediate overlap case of sensitivities in the frog skin photoreceptor system, which is similar to that of the moth *Ascalaphus* with Rhodopsin L_max_ = 354 nm and Metarhodopsin L_max_ = 480 (Hamdorf et al., 1973), a two - color capability is maintained and the risk of complete adaptation minimal in the natural environment, where yellow light is more plentiful than violet or shorter wavelength light.

Elaborating on these ideas, the double effect of any wavelength in the overlap region produces distorted measurements. When measuring excitation sensitivities, the wavelengths in the overlap region should give responses larger than the actual sensitivity yields because of their simultaneous photo - recovery effects. Conversely, when measuring spectral characteristics of recovery, overlap wavelengths will give smaller responses than the actual sensitivity would give because of simultaneous adaptation.

We have compared our spectral sensitivity curves with Dartnall’s standard curve (nomogram) for the absorption spectrum of frog rhodopsin (Dartnall, 1953). This was done on a frequency basis as the shape of absorption curves becomes independent of wavelength on the frequency scale (Wald, 1965). The spectral sensitivity curve for photo - recovery is somewhat narrower on the short wavelength side, while the photo - excitatory curve is significantly broader than the nomogram on the long wavelength side (see Fig. 8). This agreement of the nomogram with the spectral sensitivity curves suggests that the comparison may be useful. The implication for the spectral sensitivity curves — taking the nomogram curve as the real curve — is that the curves are changed as would be predicted by the wavelength overlap.

**FIGURE 8.**
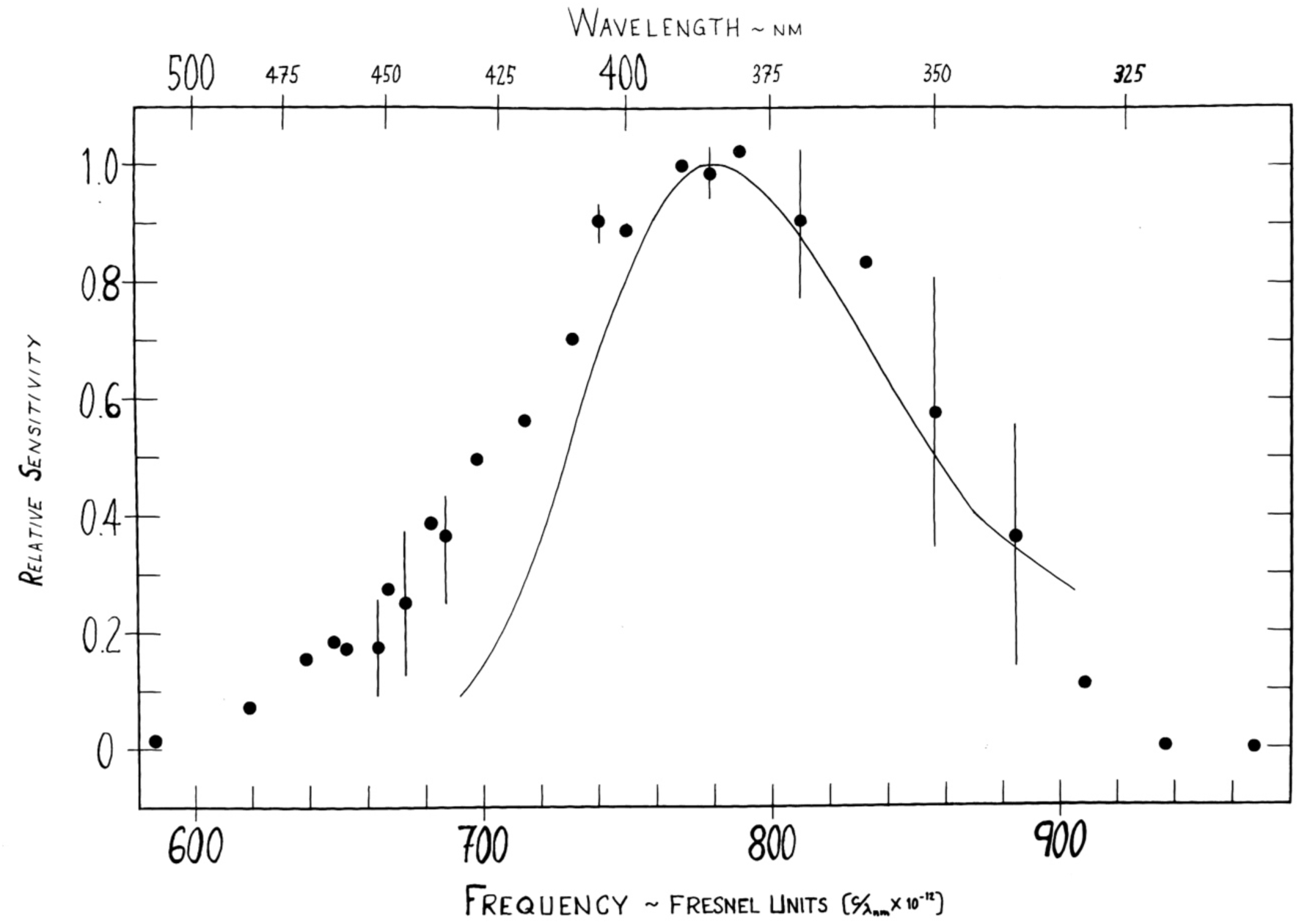
Spectral sensitivity of excitation compared with the Dartnall nomogram. The solid line is the absorption spectrum of frog rhodopsin (Dartnall, 1953) translated on the frequency scale so that it has its maximum at 385 nm. The filled circles are from the same data as used for **Figure 7** for the spectral sensitivity of excitation. If the Dartnall nomogram is a good approximation of the real spectral sensitivity curve for excitation (corrected for overlap with the recovery spectral sensitivity), then the predicted extra sensitivity measured in the overlap region (low frequency side) is the case. The error bars, again, show the range of measurements at the particular wavelength, where the circle is placed at the mean of the measurements.

## ELECTODERMOGRAM MECHANISM

Becker (1970) proposed that the skin glands mediate the EDG. Although he did not present convincing evidence for the skin gland mechanism, it is nonetheless plausible. The 50 mv reversal potential for the EDG (see Fig. 4) corresponds to the reversal potential for skin nerve stimulation where the glands are clearly implicated, in that one can see their discharge with the naked eye. Occasionally we have observed that the skin glands have discharged during the course of an experiment with light stimulation and without nerve stimulation. And, Becker (1970) found that the EDG is highly dependent on chloride in the outside bathing solution (with the absence of chloride, the EDG quickly disappears), which supports the hypothesis that the glands discharge chloride to produce the EDG, and cannot continue to do so unless they can resorb chloride from the outside. The differences in time course between the light - evoked EDG and the variation of skin potential with skin nerve stimulation may result from the use of supra - maximal nerve stimulation; when we repeated nerve stimulation at lower intensity, the correlation between the two skin responses became much closer (Rayport, 1975).

## CONCLUDING REMARKS

Skin photoreceptors in the leopard frog may be implicated as a missing link in the development of vision in vertebrates. The systematology of frog skin photoreceptors corresponds remarkably to that of invertebrate photoreceptors found in insects. The photoreversability of the system points to a sophistication beyond that of a simple pigment transformation. Various old behavioral studies suggest that skin photoreceptors may have a sensory function for the frog.

